# Screening great ape museum specimens for DNA viruses

**DOI:** 10.1101/2024.04.25.591107

**Authors:** Michelle Hämmerle, Meriam Guellil, Lovro Trgovec-Greif, Olivia Cheronet, Susanna Sawyer, Irune Ruiz-Gartzia, Esther Lizano, Aigerim Rymbekova, Pere Gelabert, Paolo Bernardi, Sojung Han, Thomas Rattei, Verena J. Schuenemann, Tomas Marques-Bonet, Katerina Guschanski, Sebastien Calvignac-Spencer, Ron Pinhasi, Martin Kuhlwilm

**Affiliations:** Department of Evolutionary Anthropology, University of Vienna, Djerassiplatz 1, 1030 Vienna, Austria; Human Evolution and Archaeological Sciences (HEAS), University of Vienna, Austria; Centre for Microbiology and Environmental Systems Science, University of Vienna, Vienna, Austria; Doctoral School of Microbiology and Environmental Systems Science, University of Vienna, Vienna, Austria; Departament de Medicina i Ciències de la Vida, Institute of Evolutionary Biology (CSIC-Universitat Pompeu Fabra), Carrer del Doctor Aiguader 88, Barcelona, 08003, Spain; Unidad de Paleobiología, ICP-CERCA, Unidad Asociada al CSIC por el IBE UPF-CSIC, Cerdanyola del Vallès, Barcelona, Spain; Institute of Evolutionary Medicine, University of Zurich, Zurich, Switzerland; Department of Environmental Sciences, University of Basel, Basel, Switzerland; Institució Catalana de Recerca i Estudis Avançats (ICREA) and Universitat Pompeu Fabra. Pg. Luís Companys 23, 08010, Barcelona, Spain; National Center for Genomic Analysis (CNAG), Baldiri i Reixac 4, 08028 Barcelona, Spain; Institute of Evolutionary Biology (UPF-CSIC), PRBB, Dr. Aiguader 88, 08003 Barcelona, Spain; Institut Català de Paleontologia Miquel Crusafont, Universitat Autònoma de Barcelona, Barcelona, Spain; Institute of Ecology and Evolution, School of Biological Sciences, University of Edinburgh, Edinburgh, UK; Department of Ecology and Genetics, Animal Ecology, Uppsala University, SE-75236 Uppsala, Sweden; Helmholtz Institute for One Health, Helmholtz-Centre for Infection Research (HZI), 17489 Greifswald, Germany; Faculty of Mathematics and Natural Sciences, University of Greifswald, 17489 Greifswald, Germany

**Keywords:** Museomics, Great Apes, Target-enrichment capture, Viruses, Hepatitis B virus

## Abstract

Natural history museum collections harbour a record of wild species from the past centuries, providing a unique opportunity to study animals as well as their infectious agents. Thousands of great ape specimens are kept in these collections, and could become an important resource for studying the evolution of DNA viruses. Their genetic material is likely to be preserved in dry museum specimens, as reported previously for monkeypox virus genomes from historical orangutan specimens. Here, we screened 209 great ape museum specimens for 99 different DNA viruses, using hybridization capture coupled with short-read high-throughput sequencing. We determined the presence of multiple viruses within this dataset from historical specimens and obtained several near-complete viral genomes. In particular, we report high-coverage (>18-fold) hepatitis B virus genomes from one gorilla and two chimpanzee individuals, which are phylogenetically placed within clades infecting the respective host species.

## Introduction

The extensive collections of fossils and specimens preserved in natural history museums document the diversity of life forms, ecosystem dynamics, and the transformative processes that have shaped our planet over millions of years. They are a critical resource for scientific research. By employing high-throughput DNA sequencing techniques tailored to ancient or historical specimens, scientists can reconstruct ancient genomes^1^ opening a new dimension in the field of museomics. Museomics is a field which has undergone rapid technical advancement, starting from using PCR products or mitochondrial DNA^4,5^, to retrieval of whole genomes^6^, while specimens preserved in liquids are another source of genomic data^7^. Spurred on by advances in the field of palaeogenomics, we can now obtain a unique picture of past genetic diversity. Beyond phylogenomic and metagenomic analysis, such material can also inform us on gene expression and regulation^8–10^. The integration of genomic technologies with traditional museum collections presents unprecedented opportunities to address consequential questions in ecology and evolutionary biology^2^. Comparing genomic data from museum specimens with modern populations allows for the reconstruction of a more comprehensive tree of life. It facilitates assessments of environmental change impacts on genetic diversity and population structure. This information is essential not only for understanding biodiversity and the forces that have shaped it in the past but also for developing effective conservation strategies to preserve genetic diversity and mitigate anthropogenic disturbances to natural ecosystems^3^.

An intriguing avenue of investigation for museomics is the study of infectious diseases^11^, as is the case for ancient DNA^12^. Natural history museum specimens have rarely been used for this purpose, but archaeological specimens have demonstrated the immense potential of ancient microbial genetic material in shedding light on the evolution and spread of infectious agents^13^. This is notably true for research on viruses with a DNA genome (DNA viruses). Hundreds to thousands of years old DNA viral genomes reconstructed from archeological specimens have unveiled key aspects of their recent evolutionary histories. For example, researchers have shown complete lineage turnover of hepatitis B viruses (HBV; family *Hepadnaviridae*) in European human populations around the Bronze Age^14,15^. Genomic pathways to increased host adaptation and virulence have also been clarified for variola viruses (*Poxviridae*) in medieval human populations^16^, for Marek’s disease virus (*Herpesviridae*) in 19^th^/20^th^ century poultry^17^, or myxoma virus (*Poxviridae*) in rabbits^18^.

In parallel, efforts geared toward a better understanding of the determinants of health in our closest relatives, the nonhuman great apes (orangutans – *Pongo pygmaeus, P. tapanuliensis* and *P. abelli*, gorillas – *Gorilla gorilla* and *G. beringei*, chimpanzees – *Pan troglodytes* – and bonobos – *Pan paniscus*), have revealed that these species host a broad range of DNA viruses. This includes enzootic viruses belonging to the families *Adenoviridae, Anelloviridae, Circoviridae, Hepadnaviridae, Herpesviridae, Papillomaviridae, Parvoviridae* and *Polyomaviridae*, as well as emerging viruses such as members of the family *Poxviridae*^19,20^. Co-phylogenetic analyses have shown that DNA viruses have often been stably associated with their hominid hosts for extensive periods of time. While co-divergence with their hosts is frequent, rare cross-species transmission events between hominids have contributed to shaping all hominid DNA viromes. For example, the herpes simplex virus 2 (*Herpesviridae*) that causes genital herpes in humans likely arose from the cross-species transmission of a virus infecting members of the gorilla lineage, several million years ago^21^. Conversely, patterns of genomic variation suggest that HBV was transmitted from humans to nonhuman great apes over the last few thousand years^15^. As we learn more about nonhuman great ape viromes, we will uncover more of the complex origins of their DNA viruses and those infecting humans.

Explorations of nonhuman great ape viromes are ongoing and should proceed in the framework of long-term studies of wild populations, but they could be efficiently complemented by analyses of the thousands of specimens kept in natural history museums around the world. Not only do these collections provide immediate access to animal tissues (as opposed to the noninvasive samples that constitute the bulk of biological sampling from modern populations), they also open a direct window on 200 years of increased anthropogenic disturbance characterized by dwindling, increasingly fragmented populations of these animals^6^. These processes have likely altered nonhuman DNA viromes significantly, and possibly resulted in the extinction of viral lineages^22^. Moreover, emerging viruses constitute a threat to wild great ape populations^20,23^, and they are often of human origin^24^. Therefore, studying the DNA viromes of nonhuman great ape museum specimens can provide insights into both extinct and contemporary viral diversity. This may also allow to monitor changes in viromes of wild populations with the potential to intervene should the populations be exposed to novel viruses (e.g. from humans or other species) they have not co-evolved with.

Here, we present high-throughput sequencing data from 209 nonhuman great ape museum specimens which were sampled and analysed in search for DNA viruses. These specimens cover all nonhuman great ape species, most subspecies and large parts of their recent geographical distribution. A majority originated in the wild, but our sample set also includes captive individuals from European zoos (13%, n=28). Obtaining data from ancient specimens can be challenging due to DNA degradation and damage^25^, as well as potential contamination during storage and handling^1^. While museum specimens are younger than typical archaeological remains, high variability in DNA quality has been observed^4^, and handling in specific clean laboratory facilities, following strict protocols is necessary^26^, and was applied for this study.

In addition, we also developed and implemented an efficient customised enrichment strategy, in-solution hybridization capture with RNA baits^27^ to account for the extremely low abundance of viral genetic material. Although metagenomic detection of viruses is possible, it is particularly challenging for DNA viruses that, with a few exceptions, usually do not replicate as intensely as RNA viruses^28^. Hybridization capture offers considerable flexibility, because it allows us to multiplex baits targeting many different viruses, as well as enrich even significantly divergent targets (up to 58% divergence tolerance according to some authors^29^). However, probably many DNA viruses evolve relatively slowly^30^. Here, we designed and used a bait set that covered the genomes of 99 viral lineages representing 13 viral families, in order to perform a cost-effective screening.

## Results

### Sequencing data

The median number of raw reads per library was 1,122,746 reads, with a high variability of yield (SD=3,556,238). After adapter trimming, a median of 956,296 reads was retained, with a reduction due to the removal of adapter-multimers between 0.11 and 93.38%. The rates of duplicated reads ranged between 21.35% and 88.67%, leaving a median of 338,560 high-quality unique reads across the 214 libraries (SD=971,276; Fig. 1A).

**Figure 1.**
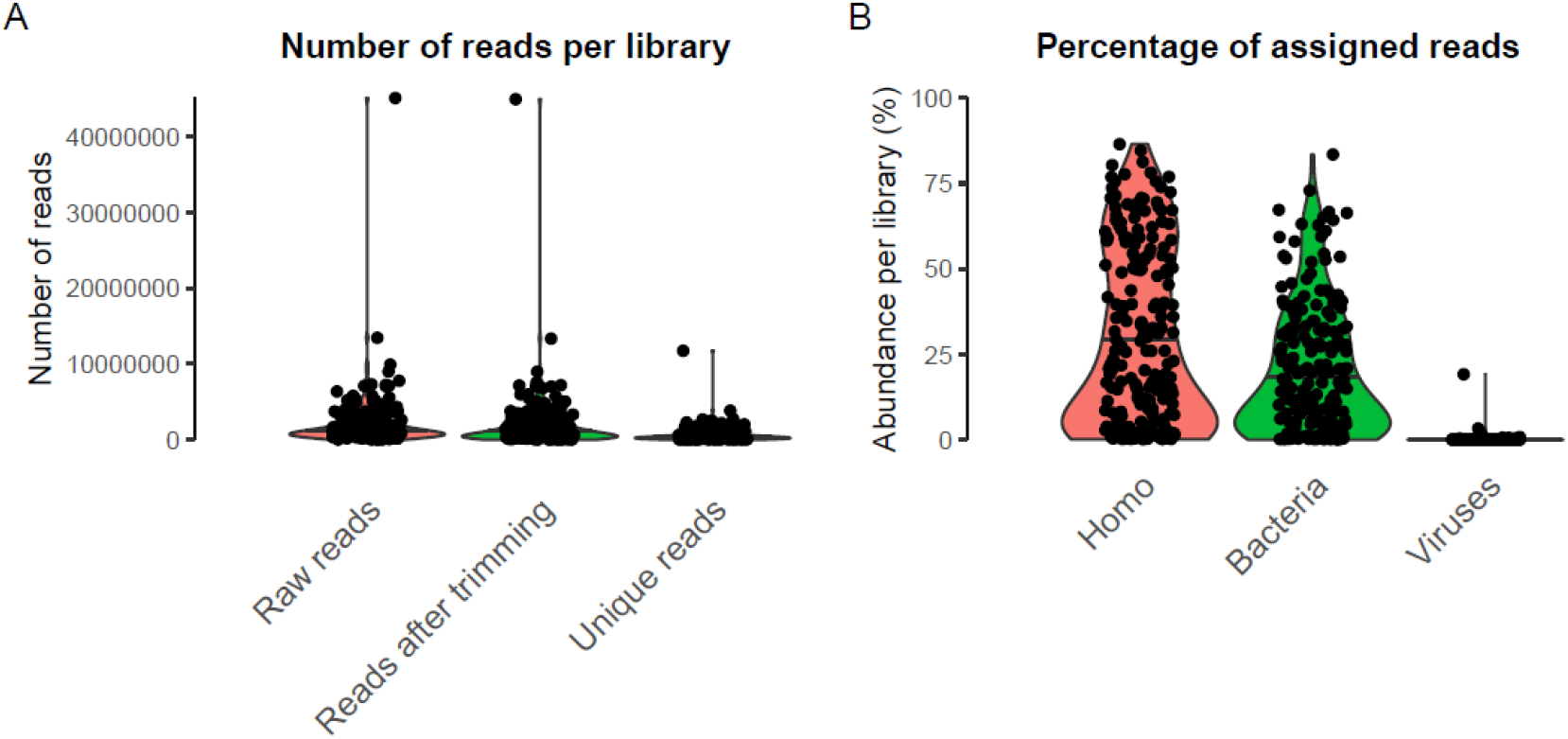
Distribution of sequencing reads per library. A) Numbers of raw reads, reads after trimming and unique reads (after BBmap clumpify) across 214 libraries. B) Percentage of reads per library assigned to *Homo*, Bacteria, or Viruses using Kraken2. We note that mapping great ape sequencing data to the human genome is commonly performed in genomic studies to avoid reference bias^31^.

The sequence length of the raw reads was 100 bp, as defined by the sequencing setup, and after filtering and trimming, the median fragment length per library ranged from 40 to 100 bp. This suggests that the extracted DNA may not have been as much fragmented as ancient DNA^1^, but likely contained a large proportion of fragments longer than 100 bp. This might be expected for museum specimens, which are younger than archaeological specimens.

### Results from virus capture

Fragments from targeted virus strains are likely absent or very rare in the libraries. Given the low expected abundance of any viral DNA, a large degree of amplification led to high duplication rates (see above). Furthermore, as we chose a lower hybridization temperature and a prolonged incubation time for the already highly amplified libraries, more unspecific binding resulted from our approach. We observed a median assignment of 21.95% to *Homo sapiens*, 13.91% to bacterial sequences and solely 0.1% to viruses (Fig. 1B). Mapping to human DNA likely reflected endogenous great ape DNA (and possibly human contamination), and bacterial DNA likely resulted from post-mortem colonization of the specimen. While many viral reads were assigned to bacteriophages (viruses infecting bacteria, not animals), many libraries contained at least one read assigned to a viral family known to infect animals (Table S3, Fig. 2A-B). Numerous libraries contained at least a small number of reads (25 or more) assigned to a viral family (Fig. 2C). A surprising amount of libraries apparently comprised poxvirus reads, which likely represented wrong assignments, likely due to high duplication in low-complexity or conserved sequence intervals. When restricting our analysis to viral taxa that were included in the capture design, we observe fewer instances (n=30) of libraries with potential virus fragments (Fig. 3), which likely reflects a more accurate picture.

**Figure 2.**
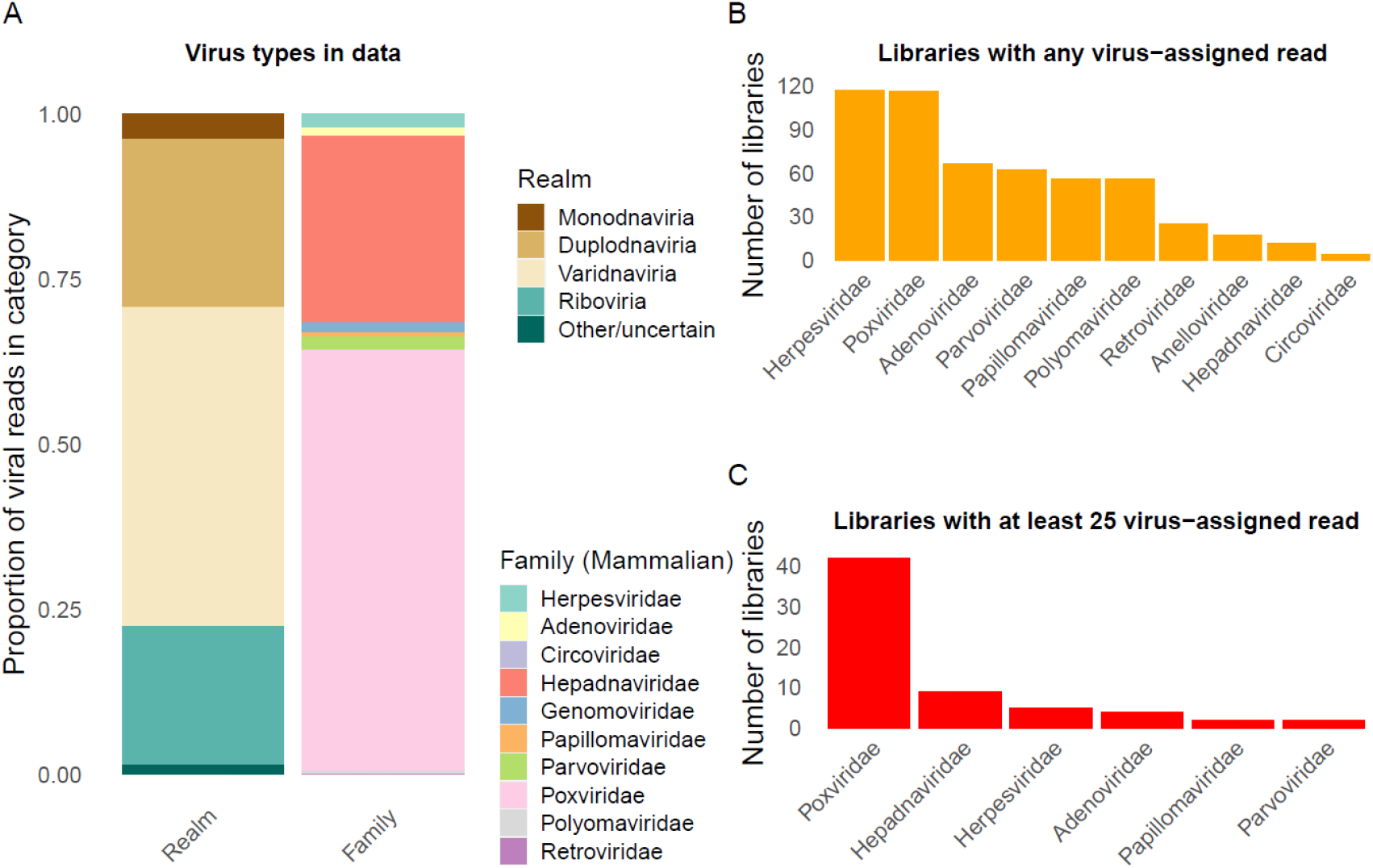
Summaries of reads assigned to different virus domains and families. A) Proportion of virus-assigned reads based on kraken2 across 214 libraries, stratified by realm and by family (for DNA viruses and *Retroviridae*). B) Number of libraries with any read assigned to virus families. C) Number of libraries with at least 25 reads assigned to virus families.

**Figure 3.**
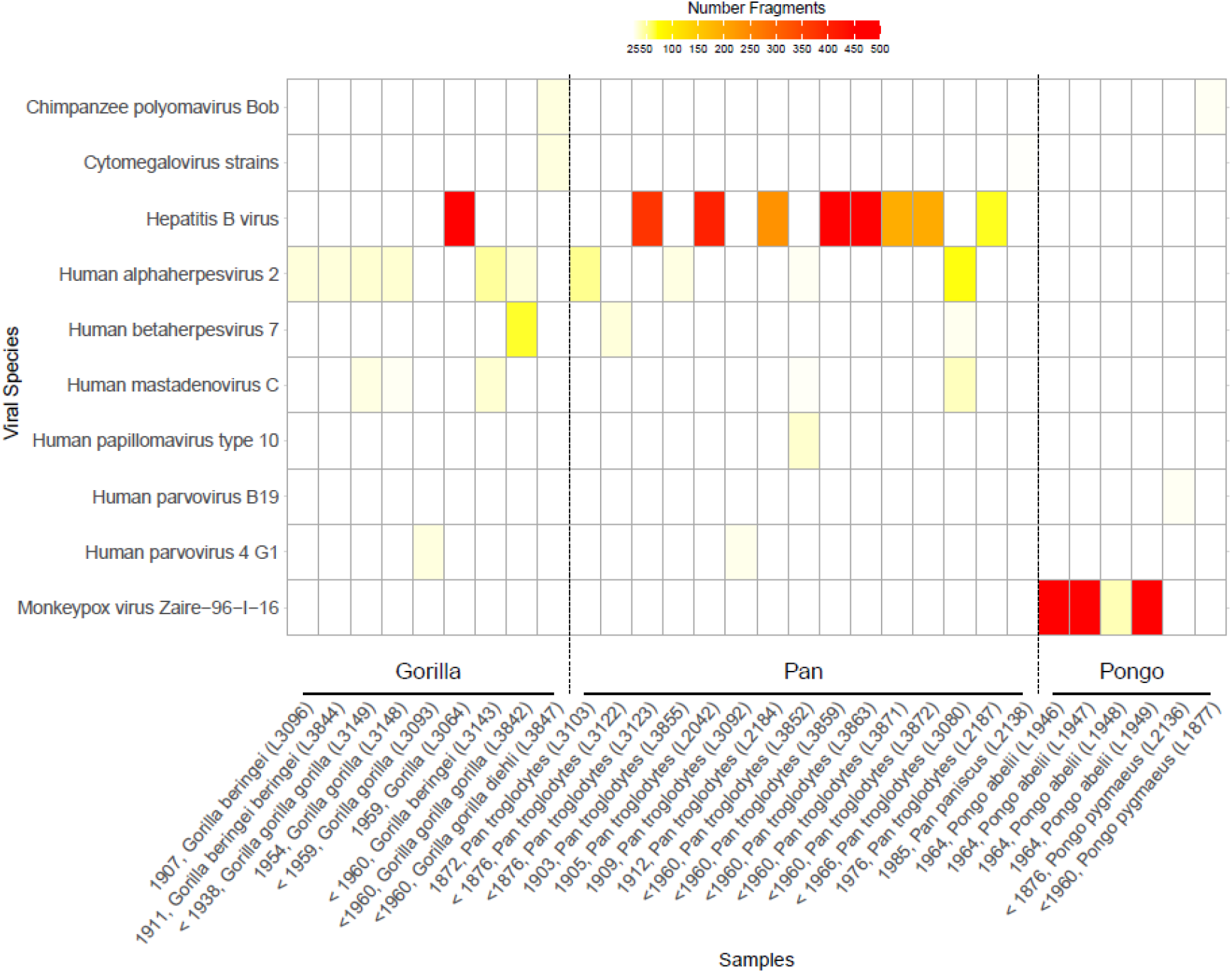
Heatmap of positive reads per viral species. The figure depicts all libraries that have at least 25 hits assigned to one of the 99 viruses in the capture kit (with merged numbers for Cytomegalovirus strains).

Henceforth, we focused on libraries with more than 500 assigned reads (based on the results obtained using kraken2), which are the most likely to reflect true viral infection. We found six such instances, with assignments either to the family *Poxviridae* (n=3; all orangutan specimens) or the family *Hepadnaviridae* (n=3; 1 gorilla and two chimpanzee specimens). The only library with more viral than bacterial or human-mapping sequences is L1949 (19.13% viral reads), obtained from an orangutan specimen. We note that further rounds of enrichment capture might have provided higher on-target coverage^1,32^, but at substantially higher costs.

### Positive specimens

To validate the results from classifying reads, we attempted to reconstruct the corresponding viral genomes. Using reference-based mapping, we managed to obtain low-coverage monkeypox virus (MPXV; family *Poxviridae*) genomes from three libraries (up to 3.6-fold coverage on the reference genome sequence KJ642614) (Table 1). Further investigation of these specimens, including deeper sequencing to obtain high-coverage genomes, was published in another study^33^, where we present their origin from a zoo outbreak in 1965. Similar efforts aimed at assembling HBV genomes from the three most promising HBV-positive specimens resulted in one high-coverage genome from a wild gorilla, and two from wild chimpanzees (Table 1), sufficient for generating consensus sequences (Methods).We could place these HBV genomes in a phylogenetic tree, verifying the placement of these strains in corresponding chimpanzee and gorilla clades (Fig. 4).

**Table 1.** Statistics for virus genomes identified in this study. Mapping metrics for the six viral genomes discovered in this study, including percentage of the reference genome covered by at least 1, 5 or 10 reads. Mapped reads = unique reads with MQ (mapping quality) >30. DOC = Depth of coverage. ED = Edit distance.

| Library ID (genus) | Individual ID (museum) | Reference | Mapped Reads | Mean ED | Mean DOC | 1X (%) | 5X (%) | 10X (%) |
| --- | --- | --- | --- | --- | --- | --- | --- | --- |
| L1946 (*Pongo*) | ZFMK_MAM1965-0547 (Bonn) | KJ642614 (monkeypox virus) | 7,931 | 0.51 | 3.83 | 93.04 | 33.25 | 2.58 |
| L1947 (*Pongo*) | ZFMK_MAM1965-0545 (Bonn) | KJ642614 (monkeypox virus) | 4,659 | 0.5 | 2.29 | 84.45 | 12.04 | 0.09 |
| L1949 (*Pongo*) | ZFMK_MAM1965-0546 (Bonn) | KJ642614 (monkeypox virus) | 2,352 | 0.61 | 1.13 | 63.83 | 1.03 | 0 |
| L3064<br>(*Gorilla*) | 96550 (Frankfurt) | FJ798097<br>(hepatitis B<br>virus) | 3,136 | 1.01 | 99.23 | 100 | 99.40 | 98.11 |
| L3859<br>(*Pan*) | ZMB_Mam_11638<br>(Berlin) | ON706349<br>(hepatitis B<br>virus) | 517 | 1.82 | 18.19 | 94.44 | 73.60 | 53.80 |
| L3863<br>(*Pan*) | ZMB_Mam_83617<br>(Berlin) | AF305327<br>(hepatitis B<br>virus) | 919 | 1.98 | 31.27 | 92.36 | 75.83 | 61.56 |

**Figure 4.**
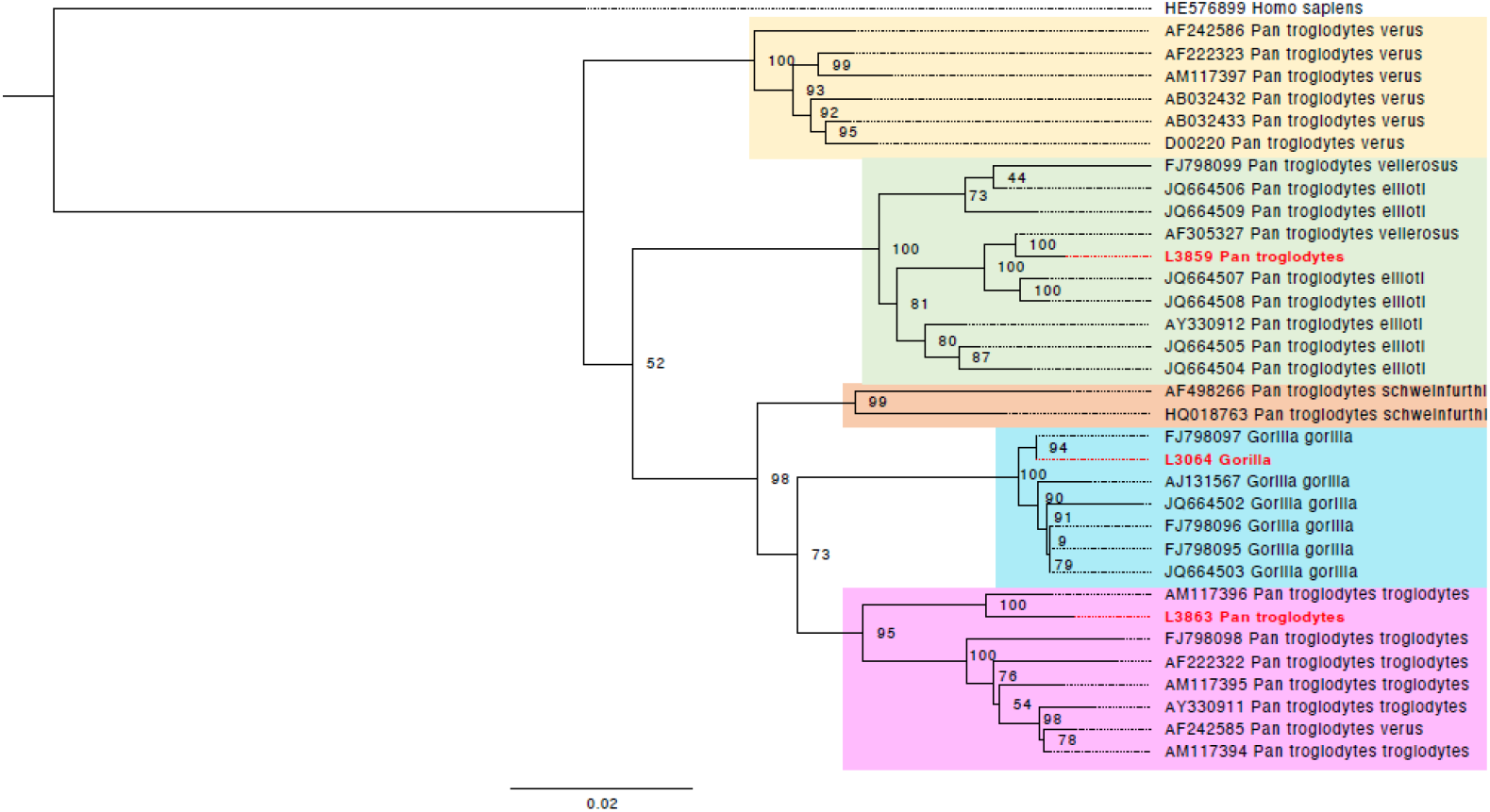
Maximum likelihood phylogeny of hepatitis B virus genomes obtained in this study, in the context of known viral diversity among great apes. The full HBV genomes were used, with a final SNPs alignment length of 4,590bp. The tree was rooted on a human hepatitis B strain, which serves as an outgroup. The colouring of the clades follows Fig. 3 in Locarnini *et al*.^35^, representing the geographical distribution of the great ape species. Samples newly sequenced in this study are marked in red.

We also attempted a *de novo* assembly strategy (Methods), focusing on the high coverage data for monkeypox virus^33^. Among the assembled viral protein fragments, 31-41% are assigned to monkeypox virus, with another 53-56% assigned to the genus or family level, some proteins to other *Poxviridae*, and indeed less than 2% to non-*Poxviridae* lineages (Table S4). We were able to recover 134 (L1946), 105 (L1947), and four (L1949) proteins with significant BLAST hits (e-value <10^-28^) and at least 90% coverage (Table S5), a substantial part of the approximately 223 open reading frames of this virus^34^.

In other samples, putative viral sequences were at a much lower abundance (Table S3).

## Discussion

In this study, we screened a total of 209 great ape museum samples with a myBaits® capture kit including 99 viral strains. We obtained six complete virus genomes, three HBV (from one gorilla and two chimpanzees), and three MPXV genomes (from three orangutans).

The high-coverage Hepatitis B genome obtained from a gorilla in this study is very similar (Fig. 4) to one sequenced from the blood serum of a wild-born western lowland gorilla from southern Cameroon^36^. The individual in our study is also a gorilla (without subspecies identification) from Cameroon, sampled more than 40 years earlier, suggesting a continuity of this lineage in the wild.

The two other HBV genomes are from chimpanzees and have a higher edit distance to the reference genomes used, indicating a deeper divergence between the genomes sequenced here, and these closest identified reference genomes. The HBV genome AM117396 from Conkouati-Douli National Park, Congo^37^ is most closely related to the HBV genome reported here for a chimpanzee from central Cameroon (Sanaga). The HBV strain AF305327 from a wild-born Nigeria-Cameroon chimpanzee^38^ is the closest match to a genome obtained from a chimpanzee individual assigned to Angola, possibly not representing the correct origin of the museum specimen. Although only the human version of HBV was included in the capture kit, the great ape-infecting lineages are nested within the diversity of human HBV lineages. Hence, the gorilla and chimpanzee versions could be identified and successfully retrieved through enrichment in this study. HBV is endemic in wild gorillas^36^ with a rate of at least 11 to 30 % of past HBV infections^39^, and finding the genetic legacy of a gorilla HBV in this study demonstrates that such infections can be detected from wild individuals from museum collections.

According to the museum metadata, the orangutan specimens yielding MPXV reads were from Sumatra. However, MPXV infections in the wild have not been observed outside of Africa. A search in the museum archive yielded a letter from the wildlife trader, stating that those were zoo individuals, without specifying the zoo from which they were taken. We identified an MPXV outbreak in Rotterdam Zoo in 1964/65^40^, during which six out of nine infected orangutans died. Mappings to an MPXV genome from this outbreak revealed a close match to the ones we obtained, leading to the conclusion that we sampled orangutans that died during this outbreak. A comprehensive study, including more data, has been published separately^33^. In this study, we were able to screen a large number of samples (209 specimens), with the inclusion of all great ape species. As great apes are endangered species (critically endangered for *Gorilla spec*. and *Pongo spec*., endangered for *Pan spec*., IUCN 2024^41^), these specimens provide a unique insight into a time when human influence on their populations was less pronounced. By pooling eight to ten samples together, it was feasible to perform an effective screening for taxa of interest. However, limitations are the availability of viral strains within databases and the use of target enrichment, which limits our detection of undescribed historical strains, as well as the fact that the targeted viruses were likely absent in some libraries. Although endogenous DNA content is low for many specimens, the target enrichment strategy is efficient and cost effective at detecting and reconstructing past viral unsampled diversity, given enrichment factors of more than 200-fold^33^. Further rounds of enrichment capture may have resulted in higher on-target rates of coverage^1,42^, but this would have significantly reduced the number of samples that could be screened. However, decreasing sequencing costs may allow a more unbiased large-scale approach in the future.

We successfully determined the presence of multiple DNA viruses in this sample set, including the detection of MPXV in three orangutans, as well as HBV in two chimpanzees and one gorilla, respectively. For these specimens, the reconstruction of near-complete viral genomes was possible, and their placement in the phylogenetic context provides evidence that these viral strains are indeed associated with their respective host species, and related to currently circulating viral lineages. Attempting *de novo* assembly from this type of data recovered a large fraction of the viral proteins for samples with high coverage. Our work demonstrates the feasibility of viral DNA enrichment and detection from museum specimens of great apes.

## Methods

### Samples

For this project, we collected a total of 209 great ape specimens, from which 214 sequencing libraries were produced. Of these libraries, 66 were from gorillas, 84 from chimpanzees (10 of these as *Pan* sp., but most likely *Pan troglodytes* ssp.), eight from bonobos, and 56 from orangutans, including different subspecies. We note that two separate extracts and sequencing libraries were prepared for five specimens. The samples were obtained from specimens housed in European natural history museums, namely in Germany in Berlin (n = 28), Bonn (n = 28), Dresden (n = 26), Frankfurt (n = 63) and Stuttgart (n = 6), in the Czech Republic in Prague (n = 24), and in Austria in Salzburg (n = 22) and Vienna (n = 12). Approximately 92% of libraries were obtained from teeth (n = 196), 17 libraries from soft tissues (Table S1), and two specimens were phalanges. According to the museum metadata, the oldest specimen was from 1838, and the most recent (a captive individual) was from 2014. Some individuals were held in captivity, mostly in zoos, whereas others were wild-caught. More detailed information concerning the individuals and the libraries can be found in Table S1.

The museum identifiers of the specimens are as follows: Senckenberg Forschungsinstitut und Naturmuseum Frankfurt/M.: 10325, 1110, 1111, 1112, 1113, 1114, 1115, 1118, 1119, 1120, 1121, 1126, 1132, 1134, 1576, 1579, 15792, 15817, 16180, 17826, 17961, 24510, 2495, 2538, 2638, 2639, 2654, 3221, 4103, 4104, 4106, 4107, 4108, 4109, 45713, 5277, 5532, 59140, 59147, 59158, 59296, 59297, 59298, 59299, 59301, 59303, 59304, 6716, 6779, 6782, 6785, 6992, 89780, 89781, 92953, 94796, 94797, 94799, 96255, 96550, 97029, 97143, ZIH9; Naturhistorisches Museum Vienna: NMW 1779, NMW 20516, NMW 25124, NMW 3081, NMW 3105/ST 663, NMW 3106, NMW 3107, NMW 3111/ST 665, NMW 3119, NMW 3948, NMW 7136, NMW 793/ST 1647; Přírodovědecké muzeum Prague: NMP 09605, NMP 10588, NMP 10784, NMP 22891, NMP 22892, NMP 22893, NMP 23283, NMP 23284, NMP 23295, NMP 23296, NMP 23297, NMP 24474, NMP 24475, NMP 46815, NMP 46816, NMP 46816-b, NMP 47007, NMP 47656, NMP 49711, NMP 50432, NMP 94205, NMP 94564, NMP 94957, NMP 95098; Senckenberg Naturhistorische Sammlungen Dresden: MTD B11877, MTD B12034, MTD B12062, MTD B12099, MTD B12101, MTD B12177, MTD B12178, MTD B1384-A.S. 1289, MTD B14244, MTD B15789, MTD B1607-A.S. 1690, MTD B247-A.S. 216, MTD B249-A.S. 214, MTD B251-A.S. 231, MTD B253-A.S. 221, MTD B266-A.S. 211, MTD B281-A.S. 239, MTD B287-A.S. 200, MTD B288-A.S. 198, MTD B3686, MTD B4188, MTD B4786, MTD B4788, MTD B4789, MTD B4793, MTD B61-A.S. 244; Museum für Naturkunde Berlin: ZMB_Mam_108652, ZMB_Mam_11637, ZMB_Mam_11638, ZMB_Mam_12799, ZMB_Mam_14644, ZMB_Mam_17011, ZMB_Mam_24838, ZMB_Mam_30755, ZMB_Mam_31617, ZMB_Mam_31621, ZMB_Mam_37523, ZMB_Mam_45130, ZMB_Mam_48173, ZMB_Mam_83519, ZMB_Mam_83522, ZMB_Mam_83547, ZMB_Mam_83606, ZMB_Mam_83607, ZMB_Mam_83617, ZMB_Mam_83642, ZMB_Mam_83643, ZMB_Mam_83647, ZMB_Mam_83648, ZMB_Mam_83653, ZMB_Mam_83675, ZMB_Mam_83681, ZMB_Mam_83682, ZMB_Mam_83685; Museum Koenig Bonn: ZFMK_MAM1938-0136, ZFMK_MAM1957-0003, ZFMK_MAM1957-0004, ZFMK_MAM1962-0131, ZFMK_MAM1963-0660, ZFMK_MAM1965-0544, ZFMK_MAM1965-0545, ZFMK_MAM1965-0546, ZFMK_MAM1965-0547, ZFMK_MAM1965-0550, ZFMK_MAM1976-0410, ZFMK_MAM1994-0482, ZFMK_MAM1997-0070, ZFMK_MAM1997-0076, ZFMK_MAM2012-0036, ZFMK_MAM2015-0479, ZFMK_MAM2019-0404, ZFMK_MAM2019-0405, ZFMK_MAM2019-0407, ZFMK_MAM2019-0408, MAM2019-0410, ZFMK_MAM2019-0415, ZFMK_MAM2019-0416, ZFMK_MAM2019-0417, ZFMK_MAM2019-0418, ZFMK_MAM2019-0419, ZFMK_MAM2019-0420, ZFMK_MAM2019-0421; Haus der Natur Salzburg: HNS-Mam-S-0073, HNS-Mam-S-0075, HNS-Mam-S-0076, HNS-Mam-S-0077, HNS-Mam-S-0078, HNS-Mam-S-0079, HNS-Mam-S-0082, HNS-Mam-S-0084, HNS-Mam-S-0085, HNS-Mam-S-0086, HNS-Mam-S-0519, HNS-Mam-S-0524, HNS-Mam-S-0525, HNS-Mam-S-0530, HNS-Mam-S-0531, HNS-Mam-S-0532, HNS-Mam-S-0533, HNS-Mam-S-0534, HNS-Mam-S-0535, HNS-Mam-S-0536, HNS-Mam-S-0550, HNS-Mam-S-0742; Staatliches Museum für Naturkunde Stuttgart: SMNS-Z-MAM-001687, SMNS-Z-MAM-001750, SMNS-Z-MAM-002012, SMNS-Z-MAM-045995, SMNS-Z-MAM-046000, SMNS-Z-MAM-048948.

### DNA extraction and library preparation

All steps from grinding to indexing except the qPCR were performed in laboratories designed and dedicated only to ancient DNA (aDNA) research while wearing protective clothing and following aDNA laboratory best practices. DNA was directly extracted from soft tissue. Bone and teeth were treated with a sandblaster and ground to bone powder using a *MixerMill* (Retsch). For each specimen, 50 mg of powder was collected. DNA was extracted using an established protocol used for aDNA^43^. Single-stranded DNA libraries were prepared^44^, followed by a clean-up with the *QIAGEN MinElute PCR Purification Kit*. We performed a quantitative PCR for calculating the cycle number in the indexing PCR. Indexing was performed in quadruplicates using *NEBNext Q5U*, followed by a clean-up using the *NucleoMag® NGS Clean-up and Size Select* kit. Indexes are listed in Table S1. We assessed quantity and quality using an *Invitrogen QubitTM 4 Fluorometer* and an *Agilent 4150 TapeStation*. The grinding step for all samples was performed in the Vienna Ancient DNA laboratory of the University of Vienna, while the DNA extraction, library and QC steps were performed for a subset of libraries (n=97, Table S1) in the ancient DNA laboratory at the Universitat Pompeu Fabra in Barcelona, following the exact same protocols and best practices.

To maximise economic efficiency, we constructed pools of 8-10 libraries. For four libraries, where a pathological condition of the individual might have influenced the specimen condition, no pooling was performed. If DNA concentration was below 5ng/μl per sample, we performed another amplification, to preserve sufficient amounts of library. If the concentration was above 25 ng/μl, a dilution was required. Between 8 and 10 libraries were pooled by equal concentration, whereby it was crucial to avoid pooling those with overlapping P5 or P7 adapters. In total, 210 libraries were pooled in 24 pools. We used the same concentration threshold for un-pooled libraries.

### Hybridization capture

As aDNA or historical DNA does not only contain host and host-associated microbiome DNA, but often an overwhelming abundance of bacterial and environmental DNA, shotgun sequencing is usually economically infeasible for viruses^45^, and target-specific approaches can help in enrichment of sequences of interest^26^. We designed a capture set containing 99 different viruses from 13 families, whereby 49 were human-infecting, 18 great ape-infecting, and the remaining were isolated from other primate species. The design was based on reference genomes available in NCBI, and commercially produced by *myBaits*^*®*^, where a BLAST search against the human reference genome was performed to exclude sequences with any hits to it; baits were designed to be 80 nucleotides long with a 2X tiling. The viruses and the NCBI reference sequence are reported in Table S2. We followed the protocol for the capture provided by the manufacturer (Version 5.03, as ordered in July 2021). Briefly, after adding the blockers and the hybridization mix with the RNA baits, the libraries were incubated for approximately 40 hours at 60°C, to increase the efficiency of the capture reaction and to allow for higher sequence deviation from the baits. DNA was eluted from the beads in 30 μl Buffer E, and the supernatant was kept. A qPCR of the capture product was performed in order to estimate the yield, and another PCR to amplify the capture product. Libraries were pooled to 20ng total DNA, and single-end sequencing was performed on the *Illumina NovaSeq 6000 (SP SR100 XP workflow)* at the Vienna BioCenter. The targeted amount of sequencing reads per library was around one million reads.

### Bioinformatic processing

Adapters were trimmed from the fastq files with trimmomatic^46^ (version 0.39), and BBmap (version 39.01) clumpify was used to remove duplicates introduced by PCR amplification^47^. To determine the metagenomic composition of the sequenced libraries, taxonomic classification via Kraken2 using the standard database was performed^48^. This database contains the strains included in the capture design. Heatmaps were plotted via a customized python3 script for the target taxa at species and genus levels.

Where Kraken2 assigned more than 500 reads to one of the reference genomes included in the capture kit, we performed a mapping with BWA^49^ (version 0.7.17), using bwa aln (with parameters “‘-n 0.04 -l 1000”‘). We performed SNP calling to the respective reference genomes presented in Table 1 using freebayes^50^, and used bcftools consensus^51^ (with parameters ‘-a “‘N”‘ --exclude ‘FILTER=“‘LOWQUAL”‘‘) to obtain consensus sequences reported in Supplementary Material SM3, with sequences for the monkeypox virus based on results with higher coverage in a separate study^33^. Mapping coverage along the reference genome was visualized using aDNA-BAMplotter^52^, and inspected individually. In case of unequal coverage along the genome, we performed a literature search for alternative genomes obtained from great apes. Summary statistics were calculated using these best reference genomes, including edit distance, mapping quality, mapping quality ratio, and the percentage for 1-, 2-, and 10-fold coverage, using a customized python3 script. Other figures were created with the R package ggplot2^53^.

For the maximum likelihood phylogeny, we used freebayes^50^ to perform SNP calling with the following flags to avoid low-quality calls (--report-monomorphic --min-alternate-count 5 -- min-coverage 5 -m 30 -F 0.9 --ploidy 1) and bcftools consensus^51^ as above to obtain consensus sequences. We included all genomes from the *Gorilla*- and *Pan*-associated genera as in Locarnini *et al*., 2021^35^. Then, a multiple sequence alignment with the 30 genomes from the previous publication and our three genomes was built via MAFFT^54^. Only positions with 90% or higher coverage were included for building the tree, which left 3,172 sites, including 695 informative ones. IQTree2^55^ was used with 1,000 nonparametric bootstrap replicates and the program chose the GTR+F+I+R2 model. Finally, the tree was formatted in Figtree v.1.4.4^56^.

An attempt for *de novo* assembly was carried out using PLASS^57^, which uses six frame translations of the sequencing reads to reconstruct proteins from a metagenome. After trimming with trimmomatic (v0.39)^46^ and removing reads smaller than 50bp, PLASS was run with the default settings except for the parameter –min-length, which was set to 20. The set of proteins obtained by PLASS was filtered to remove sequences that did not come from eukaryotic viruses, using mmseqs easytaxonomy^58,59^ with the NCBI nr database (24.10.2022)^60^ and DIAMOND (v.2.1.9)^61^ with the nr database. First, the taxonomy of the proteins was determined by easy-taxonomy and proteins being classified as coming from a virus as well as the proteins that were unclassified were additionally processed by DIAMOND blastp. Proteins that showed hits (best hits) to the nonviral proteins were filtered out. Unclassified proteins and those without hits to the nr database were mapped to vFAMs (VOGDB version 221^62^) using easy-search from mmseqs2 with the e-value cutoff 10−3. Seqtk was used for subsetting fasta files (https://github.com/lh3/seqtk). Seqkit^63^ was used to remove identical proteins from a set of proteins retrieved by PLASS.

## Supplementary Tables

Table S1. Sample metadata, including origin, year, and sample type.

Table S2. Virus strains included in the capture design, with NCBI identifiers. Table S3. Assignment of reads on the family level using kraken2.

Table S4. Number of protein sequences assigned to taxonomic units for high-coverage monkeypox data.

Table S5. Summary of BLAST hits of viral proteins to the monkeypox genome (for ORFs covered to more than 90%).

## Acknowledgements

The computational results of this work have been achieved using the Life Science Compute Cluster (LiSC) of the University of Vienna. We are grateful to the Zoologisches Forschungsmuseum A. Koenig, Leibniz-Institut zur Analyse des Biodiversitätswandels in Bonn, in particular Eva Bärmann, Jan Decher, and Christian Montermann at the Section Theriology; to Irina Ruf and Katrin Krohmann at the Mammalogy collection at Senckenberg Forschungsinstitut und Naturmuseum Frankfurt/M.; to Frank Zachos and Alexander Bibl at Naturhistorisches Museum Wien; to Stefan Merker at Zoology at Staatliches Museum für Naturkunde Stuttgart; to Petr Benda at Department of Zoology at National Museum (Natural History) Prague; to Clara Stefen and Jens Jakobitz at Mammalogie at Senckenberg Naturhistorische Sammlungen Dresden; to Robert Lindner at Haus der Natur – Museum für Natur und Technik in Salzburg; to Christiane Funk and Frieder Mayer at the mammalian collection at Museum für Naturkunde/Leibniz-Institut für Evolutions-und Biodiversitätsforschung in Berlin.

This project has been funded by the Vienna Science and Technology Fund (WWTF) [10.47379/VRG20001] and by the Austrian Science Fund (FWF) [10.55776/TAI729] to M.K. K.G. received support from the Swedish Research Council (VR) through grant 2020-03398. L. T.-G. was supported by the European Union’s Horizon 2020 research and innovation program, under the Marie Skłodowska-Curie Actions Innovative Training Networks grant agreement no. 955974 (VIROINF). T.M.-B. is supported by funding from the European Research Council (ERC) under the European Union’s Horizon 2020 research and innovation programme (grant agreement No. 864203), Grant PID2021-126004NB-100 funded by MICIU/AEI/ 10.13039/501100011033 and ERDF/EU(MICIIN/FEDER, UE) and Secretaria d’Universitats i Recerca and CERCA Programme del Departament d’Economia i Coneixement de la Generalitat de Catalunya (GRC 2021 SGR 00177). M.G. was supported by the Austrian Science Fund (FWF) [10.55776/ESP162]. S.H. was supported by the Austrian Science Fund (FWF) [10.55776/ESP546].

## Author contributions

M.K. conceived the topic of the study, supervised data analysis and wrote the manuscript. M.H. performed experiments and data analysis and wrote the manuscript. R.P. conceived the study and supervised the experimental work. M.G. supervised data analysis. S.C.-S. wrote the manuscript. S.S., E.L. and O.C. supervised experimental work. I. R.-G. performed experiments and contributed to the design of the dataset. P.B. performed experiments. L. T.-G. and A.R. analysed data. P.G., T.R., V.J.S., T.M.-B. and K.G. provided help in writing the manuscript.

## Data Availability Statement

A custom hybridization capture design for great ape DNA viruses is reported in this publication. We report the bait design for 99 virus strains of potential relevance to great apes. The final design can be found as Supplementary Material SM2, the underlying NCBI identifiers are reported in Table S2, and the NCBI sequences used in fasta format as Supplementary Material SM1. Raw sequencing data after capture for the 214 individual libraries has been uploaded to the Short Read Archive under the accession ID PRJEB75038. Consensus sequences of the discovered viruses are included as Supplementary Material SM3.

## Additional information

The authors declare no competing interests. Usage notes: Data can be reprocessed using the tools described in the Methods section, namely kraken2 for metagenomic classification, bwa for mapping to reference genomes, following the documentation in the associated repository. Custom code for processing and visualisation is available under https://github.com/admixVIE/Great-Ape-DNA-Virome.

